# Prospective isolation according to melanin pigment content of melanoma cells with heterogeneous potentials for disease propagation

**DOI:** 10.1101/2022.10.31.514484

**Authors:** Clare Fedele, Gamze Kuser-Abali, Ralph Rossi, Peinan Zhao, Jason Li, Pacman Szeto, YouFang Zhang, Nick Wong, Miles Andrews, Mark Shackleton

## Abstract

Functional variation between cancer cells (intra-tumoral heterogeneity) poses a major challenge to treating and managing cancer patients. Melanomas are typically composed of cancer cells with heterogeneous content of melanin pigment. Pigment production is a hallmark of normal melanocytic differentiation, however the functional consequences of melanin production in melanoma cells remains poorly understood owing to a lack of experimental approaches for detection of pigment in unfixed cells. Here, we describe a novel flow cytometric method for high purity separation of viable melanoma cells based on their content of melanin pigment, exploiting the infrared light scattering properties of melanin. By fluorescence-activated cell sorting, we show that melanoma cells with low-pigment content (LPCs) in culture and in patient tumors are far more abundant than high-pigment cells (HPCs), and have substantially increased potentials for colony formation in vitro and for tumor formation in vivo. RNAseq analysis revealed activation of P53 in HPCs associated with perturbed cell cycling, whereas LPCs displayed upregulation of MYC-associated transcription and activated ribosome biogenesis. The latter was reduced by topoisomerase 2 beta targeting with CX-5461, which also induced senescent HPC phenotypes and irreversible loss of clonogenic activity. These data illuminate an ‘inverted pyramid’ hierarchical model of melanoma cell propagation wherein abundant LPCs frequently renew their own malignant potential to propagate disease, but also rarely generate HPCs that lose this ability in a manner that may be promoted as an anti-melanoma strategy.

## BACKGROUND

Intratumoral heterogeneity (ITH), the presence of phenotypically, genetically, or epigenetically distinct cell sub-populations within a single tumor, is a feature of most human cancers (1, 2). Multiple models have been proposed to explain ITH, including clonal evolution based on genetic mutational instability (3, 4), reversible epigenetic ‘switching’ of cells between distinct phenotypic states (5, 6), and establishment of cellular hierarchies through irreversible epigenetic processes that resemble normal cellular differentiation (cancer stem cell [CSC] model) (7, 8). ITH has significant clinical implications if distinct cell sub-populations differ in their functional capacities to propagate malignant disease, but these differences may also illuminate therapeutic opportunities.

Melanoma is a highly heterogeneous disease (9–12), a characteristic that underlies many of the challenges in clinical melanoma management, including high propensities for development of metastasis, sometimes after prolonged dormancy, and for therapy resistance (13, 14). There is thus a critical need to understand the basis and functional consequences of ITH in melanoma. Extensive genetic ITH has been revealed in melanoma, characterized during disease progression by acquisition of sub-clones with structural genetic changes such as tetraploidization and increasing aneuploidy (15). Furthermore, the existence has been proposed within melanomas of cells with distinct phenotypic states that are largely inter-convertible and linked to degrees of melanocytic differentiation and to functional differences in cell proliferation and migration. In this ‘phenotype switching’ model (6), cells with increased features of melanocytic differentiation, such as upregulation of MITF and suppressed BRN2, display increased proliferative behavior, whereas conversion to less differentiated, MITF^lo^ states promotes melanoma cell invasion (6, 16–20). Other work showed that slow cycling cells with capacity for tumor maintenance are marked by expression of the H3K4 demethylase JARID1B, whose expression appeared plastic (21, 22).

Further studies propose unidirectional cellular hierarchies, wherein low abundance, highly tumorigenic cells generate bulk populations of less tumorigenic cells. For example, multiple groups have shown that expression of nerve growth factor receptor (NGFR/CD271/p75) can enrich for melanoma cells with tumor forming potential in patient-derived xenograft (PDX) models (23, 24). However, we and others failed to reproduce these findings in highly efficient tumorigenesis assays (12, 25). Indeed, consistent differences in tumorigenicity were not identified between melanoma cells defined by phenotypically distinct expression of a wide range of cell surface markers (12). No studies have evaluated prospectively whether degrees of canonical features of melanocytic differentiation, such as melanin production and deposition, are linked to tumorigenic potential in melanoma, although melanin was reported to inhibit melanoma metastasis (26).

Cutaneous melanins are classified as either black-brown eumelanins or yellow-red pheomelanins. Eumelanin is produced from oxidation of tyrosine by tyrosinase, forming dopaquinone, followed by cyclization to generate 5,6-dihydroxyindole and 5,6-dihydroxyindole-2-carboxylic acid, and then successive polymerization (27). Pheomelanin derives from benzothiazine units, which result from incorporation into quinones of the sulfur-containing amino acid cysteine (28, 29). It is not well understood why many melanomas retain the ability to produce melanin, and there is evidence that this retention impedes melanoma progression (26), even while it may – particularly phaeomelanin – contribute to melanomagenesis (30).

Although melanin strongly absorbs light in the ultraviolet and visible ranges (31), which is the basis for its role in protecting skin from environmental UV exposure (31–34), it is also known to exhibit light altering properties in the near infrared range (35), a property that has been used in microscopic evaluations of melanoma sections to distinguish cells in different states of melanocytic differentiation (20). We extended this approach to develop a novel method for identifying and prospectively separating melanoma cells based on their content of melanin using fluorescence-activated cell sorting (FACS), revealing intrinsic functional differences between low- (LPCs) and high-pigmented (HPCs) melanoma cells that could be exploited therapeutically. These findings refine conventional hierarchical and phenotype switching models of melanoma progression and reveal previously unappreciated complexity in the relationships between phenotypically distinct melanoma cells.

## METHODS

### Experimental Models

#### Human tumor samples

De-identified archival formalin fixed and paraffin embedded (FFPE) patient melanomas were obtained through the Melbourne Melanoma Project (MMP) and Melanoma Institute Australia (MIA). Fresh patient tumor specimens were obtained through the Victorian Cancer Biobank. Patient characteristics are listed in **Table S1**. The use of all human specimens was performed with ethical approval by the Peter MacCallum Cancer Centre Human Research Ethics Committee (HREC).

#### Mice

NOD.Cg-*Prkdc^scid^ Il2rg^tm1Wjl^/SzJ* (NSG) mice were obtained from The Jackson Laboratory. Both male and female mice were used. All mouse experiments were performed under protocols approved by the Peter MacCallum Cancer Centre Animal Ethics and Experimentation Committee (AEEC).

#### Cell lines

The B16-F10 mouse melanoma cell line and A375 and LOX-IMV1 human melanoma cell lines were obtained from ATCC and maintained in culture in DMEM supplemented with 10% fetal bovine serum (FBS) and 1% penicillin/streptomycin. The human LM-MEL-28 patient-derived cell line was a kind gift from Dr Andreas Behren at the Olivia Newton-John Cancer Research Institute and cultured in RPMI-1640 supplemented with 10% FBS and 1% penicillin/streptomycin, as described previously (36). All cell lines were maintained in 5% CO2 in a 37oC humidified cell culture incubator. Mycoplasma tests were routinely performed, and short tandem repeat (STR) profiling was conducted by the Australian Genome Research Facility (AGRF) to ensure integrity of the cell lines.

#### Xenografts

Cultured cell lines or freshly isolated patient melanoma cells were mixed with High Protein Matrigel and injected into NSG mice subcutaneously in up to three sites per mouse (back flanks and interscapular regions), as described (25). Tumors were evaluated weekly by palpation and caliper measurement.

### Cytology, Histopathology and Immunohistochemistry (IHC)

FFPE tissues were cut at 5μm thickness and mounted onto Superfrost Plus slides. Sections were dewaxed in citrolene and rehydrated through ethanol (100%, 95%, 70%) into water. For cytology studies, cells were sorted by flow cytometry directly onto Superfrost Plus slides (500-5000 cells/slide) and the drop allowed to air-dry. Slides were then immersed in dH_2_O for 5 minutes to wash away salts.

For Schmorl’s staining, tissue sections were then transferred to ferric ferricyanide solution (0.75% ferric chloride, 0.1% potassium ferricyanide in dH_2_O, prepared and filtered immediately prior to use) for 10 minutes at room temperature then washed well in tap water. Tissues were counterstained in 1% eosin for 15 seconds before being rinsed in dH_2_O, dehydrated through ethanol, cleared in citrolene and mounted in DPX mounting medium.

For Fontana-Masson (FM) staining, cells were incubated in ammoniacal silver solution (2.5% silver nitrate, 1% ammonium hydroxide in dH_2_O) in a 60°C oven for 2 hours, then with 2% gold chloride (Sigma) for 2 minutes followed by 5% sodium thiosulfate (Sigma) for 2 minutes. Nuclei were counterstained with DAPI and cells were mounted in aqueous mounting medium.

For IHC, heat-induced epitope retrieval was performed in Tris-EDTA, pH 9 solution (Dako, S2367) in a pressure cooker at 125°C for 15 minutes. Endogenous peroxidase activity was quenched in 3% H_2_O_2_ diluted in methanol for 15 minutes at room temperature. Proteins were blocked in 1% bovine serum albumin (BSA) in TBS + 0.1% Tween-20 (TBS-T) for 30 minutes at room temperature. Sections were then incubated with anti-S100 primary antibody (1:500, Dako, Z0311) overnight at 4°C in a humidified chamber, then with biotinylated anti-rabbit IgG ((goat, 1:200, Abcam, ab6720). An avidin-HRP-Biotin Complex (ABC) system (Vector Labs) was applied according to manufacturer’s instructions, and immunoreactivity revealed using AEC (Dako). Tissues were counterstained with haematoxylin then cleared and mounted as above.

For detection of cell senescence with X-galactosidase, 1.5×10^4^ cells were cultured on glass coverslips in a 24-well dish for 5 days to adhere. Cells were fixed in 2% formaldehyde/0.2% glutaraldehyde/PBS for 5 minutes then incubated in X-galactosidase staining solution (5 mM potassium ferrocyanide, 5 mM potassium ferricyanide, 150 mM NaCl, 2 mM MgCl_2_, 1 mg/mL X-galactosidase in phosphate/citrate buffer (7.16 mM citric acid, 25.5 mM NaH_2_PO_4_); all from Sigma) overnight in a non-humidified 37°C incubator. Nuclei were counterstained with DAPI and cells mounted in ProLong Gold.

Proportions of HPCs in total cell populations was determined by either light microscopy or Schmorl’s/Fontana-Masson staining and expressed as a % of total cell nuclei as determined by DAPI or haematoxylin counterstains. Cell diameters were measured in Image J (NIH) on FM stained, non-adhered cells sorted onto slides. Differences in cell diameter were calculated using an unpaired Student’s t-test.

### Preparation of Tumor Single Cell Suspensions

Fresh patient tumors and patient-derived xenografts (PDX) were dissociated using the same method, adapted from (37). Tumors were mechanically dissociated with a McIIwain tissue chopper (Mickle Laboratory Engineering) and the tumor slurry was resuspended in HBSS (without Ca2+ and Mg2+ (HBSS-/-)), Sigma) containing 200 U/mL collagenase IV (Worthington) with 50 U/mL DNase (Roche) and 1 mg/mL CaCl_2_ (10 mL of digestion media per gram of tissue), and placed in a 37°C water bath for 20 mins, with agitation and mixing every 5 mins. The resultant digest was washed with HBSS-/- and spun down at 220g for 4 mins at 4°C. The cell/tissue pellet was then gently resuspended in warmed 0.05% trypsin-EGTA (Gibco) with 200 U/mL DNase, followed by incubation at 37°C for 5 mins. More DNase (50-200 U/mL) was added when necessary to reduce cell clumping during digestion. An equal volume of chilled staining media (L15 medium (Gibco) containing 1 mg/mL BSA, 1% penicillin/streptomycin, 10 mM HEPES (pH7.4)) was added to quench the trypsin, followed by centrifugation at 220g for 4 mins at 4°C. The cell pellet was resuspended in staining media and filtered through a 40 μm cell strainer to obtain a single cell suspension.

### Flow Cytometry

For patient-derived melanomas, antibody labeling was performed for 20 minutes on ice. Cells from patient samples were stained with directly conjugated antibodies to human HLA-A, B, C (1:5, G46-2.6-FITC, BD Pharmingen), human CD45 (1:5, HI30-APC, BD Pharmingen), human CD31 (1:800, WM59-APC, eBioscience) and CD235a (glycophorin A; 1:2000, GA-R2 (HIR2)- APC, BD Pharmingen) to enable selection of HLA^+^CD45^-^CD31^-^CD235a^-^ (Lin^-^) cells (**Figure S2**). Cells from PDX tumors were stained with directly conjugated antibodies to human HLA-A, B, C (as above), mouse CD45 (1:200, 30-F11-APC, BD Pharmingen), mouse Ter119 (1:100, TER119-APC, BD Pharmingen), and mouse CD31 (1:100, 390-APC, eBioscience). DAPI (Roche) was used as a marker of cell death, and DAPI^-^ viable cells were gated for downstream analyses.

Mouse and human adherent cultured cell lines were detached in Trypsin-EDTA (Gibco) and resuspended in staining media with DAPI for viability. Endogenous melanin was detected using a customized FACSAria II (BD Biosciences) fitted with a near-infrared laser. Filters tested for melanin detection were 710 long-pass (LP), 745 LP, 750 LP, 800 LP, 900 LP, 817/25 band-pass (BP) and 800/30 BP (BD Biosciences). The 800/30 BP filter was used for all subsequent analyses. Melanin content was defined based on side-scatter (SSC) and near-infrared scatter (NIRSC).

For detection of stem cell markers cells were non-enzymatically detached from plastic in citric saline (0.135 M citric acid, 0.015 M sodium citrate in H_2_O). All antibody labelling was performed at room temperature for 30 minutes. For detection of cell surface markers in unfixed cells, live cells were incubated with fluorescently conjugated or unconjugated primary antibodies, or matched IgG controls at the same concentration. Primary antibodies were: anti-CD271-PE (Miltenyi, #120-002-227), anti-c-Kit-PE (eBioscience, #12-1179-41), anti-CD49f-FITC (BD Biosciences, #555735), anti-ABCB5 (Sigma, #HPA026975), anti-ALDH1A1 (Abcam, #ab52492), anti-L1CAM (Abcam, #ab24345), anti-CD51 (BD Biosciences, #611013), anti-A2B5 (Invitrogen, #433110) and anti-GD2 (BD Biosciences, #554272). For detection of intracellular proteins, cells were first fixed in 4% formaldehyde/PBS for 15 minutes at room temperature, then permeablized in ice cold 90% methanol/PBS for 30 minutes on ice, before incubation with primary antibodies, or IgG controls: anti-JARID1B (Abcam, #ab84883), anti-SOX10 (Abcam, #25978), anti-ABCB5 (Sigma, #HPA026975) and anti-MITF (Merck Millipore, #MABE78). Unconjugated primaries were detected using appropriate AlexaFluor secondary antibodies (ThermoFisher). Marker-positive populations were gated based on signal above background as determined by IgG controls, following laser voltage correction for increased basal background signal in HPCs.

For detection of apoptosis by flow cytometry, adherent and floating vehicle- or CX5461-treated cells were resuspended in AnnexinV binding buffer (0.14M NaCl, 0.01M HEPES, 2.5mM CaCl_2_) and 2 μL AnnexinV-FITC was added to each tube and incubated on ice for 30 minutes. Nuclei were counterstained with 5 μg/mL propidium iodide.

### *In vitro* Drug Treatments

To stimulate melanin production, 2.5×10^5^ B16-F10 cells were plated into a 6-well dish and stimulated with growth media supplemented with 10 or 20 μM forskolin (Sigma) and 100 μM 3-isobutyl-1-methylxanthine (IBMX) (Sigma), or DMSO and methanol vehicle controls, under normal culture conditions for 72 hours prior to flow cytometric analysis.

For inhibition of topoisomerase 2 beta, 2×10^5^ 28F3:B4 cells were seeded into 12-well dishes then stimulated with growth media supplemented with 1 μM CX5461 (a kind gift from Rick Pearson, Peter MacCallum Cancer Center) or NaH_2_PO_4_ vehicle for 5 days (for in vitro assays) or 10 days (for in vivo assays) under normal culture conditions, replacing fresh media with drug every 2-3 days.

### *In vitro* Clonogenicity Assays

For single cell clonogenicity assays, single LPCs or HPCs were sorted based on SSC/NIRSC characteristics into growth media in 96-well tissue culture-treated dishes, 1 cell/well. At least 48 cells/phenotype/experiment were plated. Cells were briefly centrifuged at 220g to collect at the bottom of the well, then each well was assessed by light microscopy to confirm the presence of a single cell and the phenotype (LPC or HPC) of each cell, and imaged. Any cell that did not match the expected phenotype of the sorted population was excluded from further analysis. Cells were then cultured for 21-28 days under standard culture conditions, changing media every 5-7 days. At the conclusion of the experiment, each well was re-assessed by light microscopy for colony formation and phenotype and colonies were re-imaged. Differences in clonogenicity were calculated using a 2×2 contingency table with Fisher’s exact test.

For limiting dilution clonogenicity assays using extreme limiting dilution analysis (ELDA) (38), vehicle- or CX5461-treated cells were detached and re-plated into growth media in 24-well plates at 100, 10, 5 and 2 cells/well in six replicates. Colonies were allowed to form under standard growth conditions for 28 days, changing the media every 5-7 days. At the conclusion of the experiment, colonies were fixed in 100% methanol for 45 minutes then stained in 0.1% crystal violet for 45 minutes.

### *In vivo* Tumorigenicity Assays

Purified LP or HPCs, or vehicle- or CX5461-treated cells, were counted, mixed with 25% High Concentration Matrigel (BD Biosciences) and injected subcutaneously into NSG mice at limiting dilutions (1000, 100, 10 cells/injection). At experimental endpoint, tumors were excised and necropsies performed on mice to assess macroscopic metastasis in visceral organs. Tumor latencies were calculated as time from injection until tumors first became palpable. Tumor growth rates were calculated as maximum mm diameter/week from day of first palpation to experimental endpoint.

### RNA Sequencing and Gene Expression Analysis

For transcriptomic analysis, total cell RNA was extracted from FACS-purified LP or HP cell populations from B16-F10 murine or LM-MEL-28 human melanoma cells using PureLink RNA Mini kits (ThermoFisher), including on-column DNase treatment. RNA was quantitated using Qubit RNA HS (Thermo Fisher Scientific) and 100 ng total RNA input was used for library preparation according to standard protocols (QuantSeq 3’ mRNA-Seq FWD, Lexogen). Indexed libraries were pooled and sequenced on a NextSeq 500 sequencer (Illumina). 5-15 million single-end 75bp reads were generated per sample. Fastq files for B16-F10 and LM-MEL-28 were aligned to GRCm38 and GRCh37 respectively. Raw gene count matrices were generated using featureCounts from the Subread package (http://subread.sourceforge.net). Differential expression (DE) analyses were performed using limma-voom (v.3.48.3). Significant DE genes (FDR<0.05) were used for enrichment analyses to identify GO biological process and Hallmark gene sets, performed using clusterProfiler (v3.18.1) (39). Over-representation analysis was conducted based on hypergeometric distribution, and p values adjusted by the Benjamini-Hochberg (BH) procedure (40). Significantly enriched pathways were defined as those with an adjusted p value < 0.05. The COSMIC cancer gene census (https://cancer.sanger.ac.uk/census) was referenced to identify oncogenes and tumor suppressor genes. RNAseq data were deposited in Gene Expression Omnibus (GEO) (accession number GSE214826 and GSE190252).

### Software and Statistical Analysis

Flow cytometry data were analyzed and quantitated using the Weasel v3.0.2 flow cytometry display and analysis program (Walter and Eliza Hall Institute). Unless otherwise stated, statistical analyses were performed using R (v.4.0.4) for RNAseq data and GraphPad Prism 7 software for other data. For statistical analyses, p<0.05 was the cut-off for significance, except for GSEA where FDR q<0.25 was used. Sample sizes and statistical details are provided throughout the text and in figure legends.

## RESULTS

### Heterogeneity of cellular melanin pigment content is common in pigmented melanomas, melanoma cell lines and patient-derived xenografts

To examine heterogeneity of cellular melanin pigment content amongst melanoma cells, we first performed Schmorl’s staining in a cohort of 51 stage I/II primary melanomas and 65 stage III/IV melanoma metastases obtained from the Melanoma Research Victoria cohort. Of the 116 melanomas stained, 93 (80%) were pigmented, of which 85 (91%) exhibited heterogeneous melanoma cell pigmentation. Cells with no or only a faint blush of pigment were called ‘low pigment cells’ (LPCs) and cells with at least moderate levels of pigment were called ‘high pigment cells’ (HPCs) (**Figure 1A**, **Figure S1A**), and LPCs were confirmed to be melanoma cells by S100 labelling (**Figure S1B**). Although metastatic melanomas were more likely than primary cutaneous melanomas to contain only LPCs, consistent with previous observations (41), pigmented melanomas from different disease stages displayed similarly abundant IYH of pigment deposition; cells in 44 of 48 (92%) pigmented primary melanomas and in 41 of 45 (91%) pigmented metastatic melanomas were heterogeneously pigmented.

**Figure 1.**
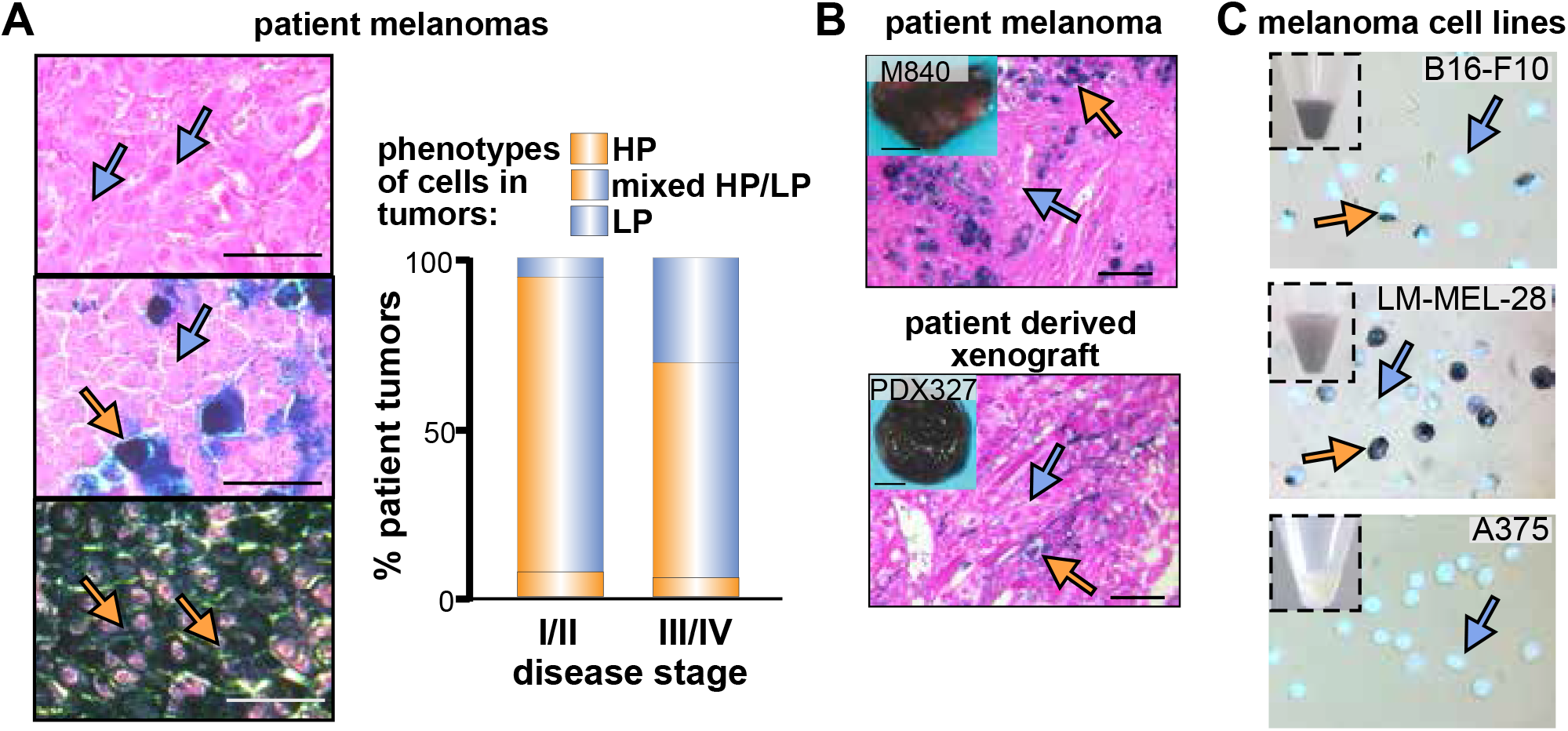
Most melanotic melanomas are heterogeneously pigmented. **A)** Pigmentation patterns in patient melanomas. Left: Schmorl’s staining of FFPE sections of patient melanomas showing the patterns of cell pigmentation seen; blue arrows: low pigment cells; orange arrows: high pigment cells; scale bars: 50μm. Right: quantitation of the proportion of patient melanoma that contained high pigment (HP) cells only, low pigment (LP) cells only, and heterogenous (mixed) HP and LP cells. **B)** Schmorl’s staining of pigmented patient (top) and patient-derived xenograft (bottom) melanomas from patient M840; bars: 50 μm. **C)** Fontana-Masson (FM) staining of cytospins of macroscopically amelanotic (A375) and melanotic (B16-F10 and LM-MEL-28) melanoma cell lines (insets show pelleted cells).

To determine if pigment ITH is maintained during patient-derived xenografting (PDX) we injected into immunocompromised NSG mice FACS-purified, unselected melanoma cells derived from pigmented human melanoma tumors obtained fresh from surgery (12, 25, 37). PDX tumors recapitulated the pigment phenotypes of parental tumors, forming overtly pigmented tumors comprised of heterogeneously pigmented cells (**Figure 1B**). We also examined pigment heterogeneity by Fontana-Masson (FM) staining of cytospins of pigmented melanoma cell lines, finding heterogeneous mixtures of LPCs and HPCs in B16-F10 mouse melanoma cells and in patient-derived human LM-MEL-28 cells (36). In contrast, amelanotic human A375 melanomas cells comprised only LPCs (**Figure 1C**). Interestingly, the LPC state was more abundant than the HPC state in both pigmented B16-F10 and LM-MEL-28 lines. Thus, heterogeneity in cell melanin pigment content is a feature of macroscopically pigmented patient melanomas, PDX melanomas and melanoma cell lines.

### Melanoma cells can be separated based on their melanin pigment content by fluorescence-activated cell sorting

As melanin pigment production in normal melanocytes is a feature of terminal differentiation, we reasoned that pigment production in melanoma cells may be linked to differences in tumorigenicity/clonogenicity. However, prospective testing of this has not been possible due to an inability to separate melanoma cells based on their content of melanin pigment. As melanin pigment is packaged within specialized organelles called melanosomes (42), we hypothesized that HPCs would possess increased melanosome number and density, compared to LPCs, and that these would be detectable by flow cytometry as side scatter (SSC) of 488 nm light. Consistent with this, SSC^hi^ cells were enriched in pigmented B16-F10 cells but not in amelanotic A375 cells (**Figure 2A**).

**Figure 2.**
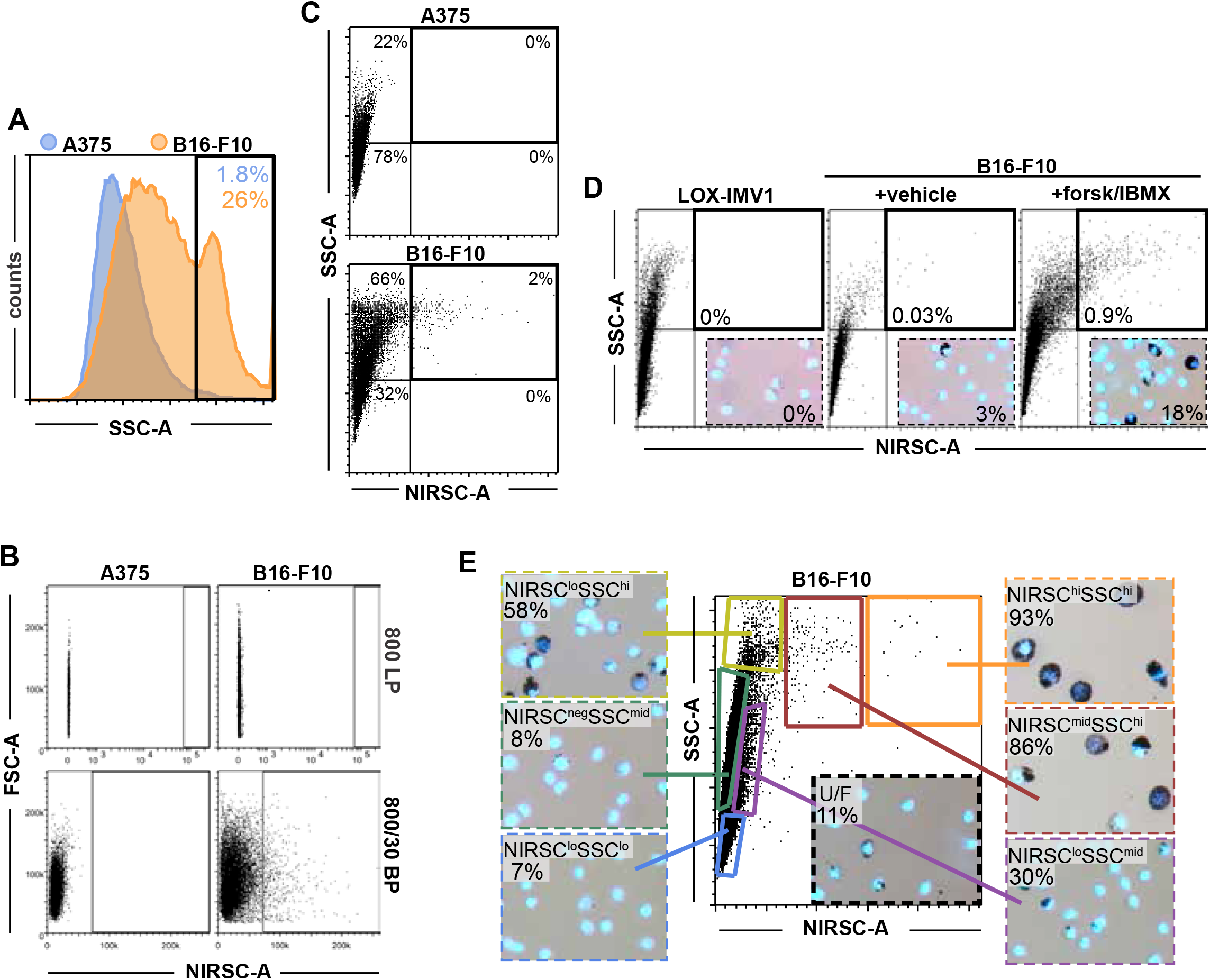
Light scattering properties of differentially pigmented melanoma cells. **A)** Flow cytometric analysis of side scatter (SSC) in amelanotic A375 and heterogeneously melanotic B16-F10 cells. **B)** Flow cytometric detection of signal following excitation of A375 or B16-F10 cells with near-infrared (792nm) wavelengths and detection with either 800 long-pass (LP) or 800/30 band-pass (BP) filters. **C)** Flow cytometric detection of SSChiNIRSChi cells in melanotic B16-F10 but not amelanotic A375 cells. **D)** Flow cytometric analysis of SSC and NIRSC in amelanotic LOX-IMV-1 cells or B16-F10 cells treated for 72 hours with 20μM Forskolin (Forsk) + 100μM IBMX, or control vehicle. % of SSChiNIRSChi cells is indicated. Inset: FM staining for melanin and % FM+ cells. **E)** Gating of B16-F10 cells based on SSC/NIRSC characteristics. % FM+ HPC cells are indicated in unfractionated (U/F) cells or in each sorted population.

To augment the distinction of LPCs and HPCs, and as melanin was reported to possess intrinsic fluorescence when stimulated by near-infrared (NIR) light (20, 43), we excited melanoma cells with 792 nm light. However, we could not detect emissions by flow cytometry using either 800 nm or 900 nm long-pass detection filters (**Figure 2B**, **Figure S2**). Instead, a population of NIR^hi^ cells was detected in B16-F10 cells, but not A375 cells, using either 800/30 band-pass (BP) (**Figure 2B**) or 710-750 nm long-pass filters (**Figure S2**), each of which detects 792 nm excitation light wavelengths.

As these detection patterns were more consistent with NIR light scattering rather than classical excitation/emission fluorescence, we termed this “near-infrared scattering” (NIRSC) and hypothesized that cellular pigmentation would be proportional to both SSC and NIRSC in dual-colour flow cytometry. Accordingly, a population of SSC^hi^NIRSC^hi^ cells was detectable in B16-F10 cells but not in amelanotic A375 cells (**Figure 2C**), and the SSC^hi^NIRSC^hi^ population increased with addition to cultures of pigment-inducing compounds forskolin and 3-isobutyl-1-methylxanthine (IBMX), together with increasing FM positive cells (**Figure 2D**, **Figure S3A**).

To confirm that SSC^lo^NIRSC^lo^ and SSC^hi^NIRSC^hi^ cells enrich for LPCs and HPCs, respectively, we conducted fluorescence-activated cell sorting (FACS) on melanoma cells isolated from a macroscopically pigmented PDX melanoma that exhibited pigment cell heterogeneity by FM staining (**Figure S3B**). SSC^lo^NIRSC^lo^ cells were devoid of pigment (**Figure S3Bi**), while increasing stringency of SSC^hi^NIRSC^hi^ gating increased the purity of HPCs (31-89%) (**Figure S3Bi-iii**). A similar approach in B16-F10 cells demonstrated that increasingly restricted gating of SSC^lo^NIRSC^lo^ and SSC^hi^NIRSC^hi^ populations progressively enriched for LPCs and HPCs, respectively (**Figure 2E**). We thus describe a novel method for prospectively purifying unfixed and viable cells according to their content of melanin pigment.

### LPCs have increased colony forming potential *in vitro*

To test whether LPCs are more clonogenic than HPCs, we availed SSC/NIRSC-based cell sorting to compare fresh and prospectively isolated cells in side-by-side assays. As cell size has been linked to clonogenic potential (44), we first noticed that isolated LPCs were consistently smaller than HPCs in B16-F10 and LM-MEL-28 cells, as well as cells from a dissociated PDX melanoma (mean diameter 15.6 μm and 23.7 μm, respectively) (**Figure 3A**). We next sorted maximally enriched, single LPCs and HPCs directly into 96-well plates and confirmed the presence of single cells in each well by direct light microscopic visualization (**Figure 3Bi-ii**). Cells were cultured and a resultant colony was defined as any well containing ≥2 cells proximal to each other, indicating at least one cell division. In B16-F10 cells, a ~2-fold higher number of single LPCs than HPCs formed colonies (45% and 23%, respectively) (**Figure 3Ci**). Approximately 20% of LPCs and HPCs displayed limited proliferative activity, forming colonies of <10 cells, equivalent to 1-3 cell divisions. In contrast, cells that formed colonies of ≥10 cells, and particularly of >100 cells, were almost exclusively LPCs. Similar results were obtained for LM-MEL-28 cells (**Figure 3Cii**) and for melanoma cells dissociated and purified from PDX melanoma M442 (**Figure 3D**).

**Figure 3.**
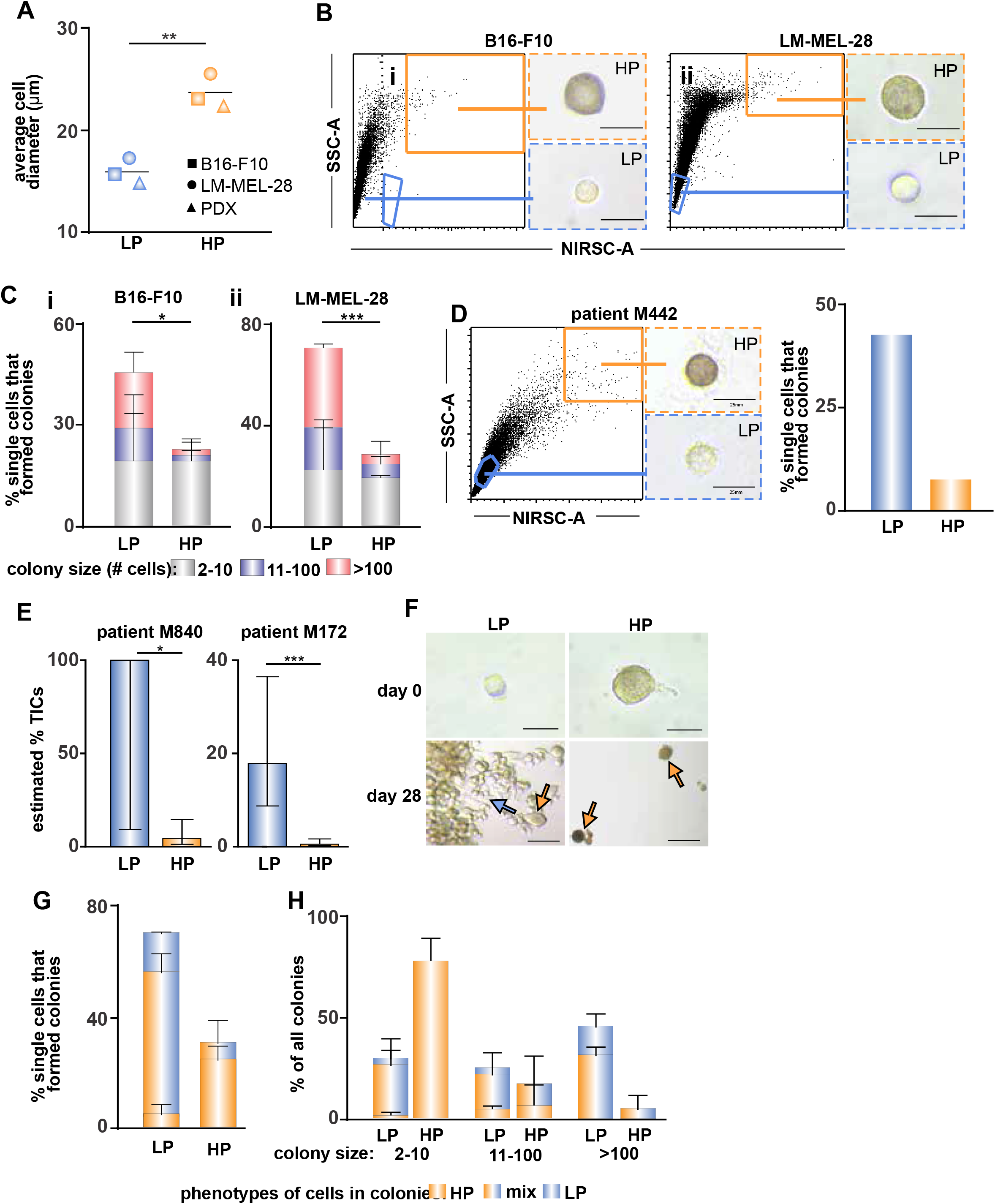
LP cells have increased clonogenic and tumorigenic potentials. **A)** Average diameters of LPCs and HPCs sorted from B16-F10 cells, LM-MEL-28 cells or a pigmented PDX melanoma. n≥30 cells were measured per phenotype per line. **p<0.005, unpaired student t-test. **B)** Single HP or LP cells from i) B16-F10 or ii) LM-MEL-28 were sorted by flow cytometry based on SSC/NIRSC (1 cell/well). Each sorted cell was assessed for pigment by light microscopy, then maintained for 21-28 days in culture. **C)** Quantitation of colony (>2 cells) formation and the number of cells per colony for i) B16-F10 or ii) LM-MEL-28 cells. Bars: averages ± SD from 2 independent experiments. n ≥ 17 single cells/phenotype/cell line/experiment. *p<0.01 and ***p < 0.001, t-tests. **D)** Single HP or LP cells from a pigmented PDX melanoma from patient M442 were sorted into single wells in vitro and cultured for 15 days. At endpoint, colony formation (>2 cells) was assessed. Bars: 25 μm. **E)** Extreme Limiting Dilution (ELDA) analysis of tumor-initiating cell (TIC) frequencies of LP and HP cells from 2 PDX melanomas. Bars: estimated % TICS ± upper and lower limits. Injections/phenotype: M840: 2x 1000 cells, 2-3x 100 cells and 6x 10 cells, and M172: 6x 100 cells and 12x 10 cells. **F)** Single, sorted LP or HP LM-MEL-28 cells were assessed on day 0 for pigment by light microscopy (upper panel), then cultured for 28 days. At experimental endpoint, resultant colonies were assessed by light microscopy (lower panels) for LP (blue arrows) and HP (orange arrows) cells. Bars: 25 μm (upper) and 100 μm (lower). **G)** Quantitation of colony phenotypes grown from single LP or HP cells. Bars: averages ± SD from 2 independent experiments. n ≥ 17 single cells/phenotype/experiment. **H)** Quantitation of colony phenotypes stratified by colony size. Bars: averages ± SD from 2 independent experiments. n ≥ 7 (HP) and n = 32 (LP) colonies/experiment.

### LPCs have increased tumor forming potential *in vivo*

To test the tumorigenic potential of LPCs and HPCs *in vivo,* we established PDX melanomas in NSG mice (M840 and M172) and used assays that had previously permitted very efficient tumor formation by melanoma cells regardless of their cell surface marker phenotype (12, 37). Purified LPCs and HPCs from these PDX melanomas (**Figure S4A**) were re-injected into NSG mice in decreasing numbers and the frequency of tumor-initiating cells in each phenotype calculated from rates of consequent tumor formation (38). Like colony formation, cells with tumorigenic potential were more abundant in LPCs than HPCs isolated from either melanoma (**Figure 3E**). Tumors that grew from HPCs had longer latencies than tumors grown from LPCs (**Figure S4B**), but similar tumor growth rates were observed once tumors became palpable from either phenotype of injected cells (**Figure S4C**). Furthermore, although *MITF*^lo^/undifferentiated melanoma cells were found to be more invasive than *MITF*^hi^/differentiated melanoma cells (6, 45), mice with subcutaneous tumors grown from either LPCs or HPCs developed widespread metastases (**Figure S4D)**. These data implicate deposition of melanin pigment in melanoma cells as a marker of decreased tumorigenic potential, even when tumorigenicity is tested in highly permissive assays.

### LPCs but not HPCs typically generate heterogeneously pigmented clonal colonies

We also assessed the pigmentation phenotypes of clonal colonies derived from LPCs and HPCs. For both B16-F10 and LM-MEL-28 cells, the mostly small colonies formed by single HPCs typically comprised only HPCs that could not be long-term expanded. On the other hand, most colonies derived from single LPCs contained heterogeneously pigmented cells, recapitulating the pigment heterogeneity of their parental cultures (**Figure 3F-H**, **Figure S4E-G**). Long-term (>2 months) expansion of four clonal LPC-derived colonies resulted in re-equilibration of pigment heterogeneity (**Figure S4G**). Thus, the ability to recapitulate colony heterogeneity was enriched in LPCs.

In vivo, PDX tumors formed from either LPCs or HPCs both contained heterogeneously pigmented melanoma cells pigment (not shown). This could reflect a greater degree of plasticity within the HPC state in vivo than seen in vitro, however it cannot be ruled out that PDX tumors grown from HPCs may have in fact been derived from rare LPCs remaining in highly enriched but impure HPC suspensions after sorting.

### Clonal passaging of LPCs and HPCs

As some LPC clones were expandable long-term and able to recapitulate the phenotypic heterogeneity of parent populations, we hypothesized that LPCs were enriched for self-renewing cells, in which case daughter LPCs would maintain the potentials for clonogenicity and multipotency of parent LPCs. To test this, we expanded a single LPC-derived, heterogeneously pigmented colony from the LM-MEL-28 line (LM-MEL-28 clone F3), and performed a second round of single cell plating, forming a second-generation clonal colony (28:F3:B4). 28:F3:B4 cultures were heterogeneously pigmented, like the parental LM-MEL-28 and F3 clonal lines (**Figure 4A**). In single cell in vitro assays, flow cytometrically purified 28:F3:B4-derived LPCs were enriched for clonogenic cells compared with HPCs (**Figure 4B-C**), as observed for parental lines. Single 28F3:B4 LPCs generated colonies (**Figure 4D**) that were more likely than HPC-derived clonal colonies to be large (>100 cells) and heterogeneously pigmented (**Figure 4E**). In contrast, single 28:F3:B4 HPCs-generated colonies were more likely to be homogenously pigmented and smaller (**Figure 4D-E**). Interestingly, when HPCs derived colonies of >10 cells, these colonies were comprised of heterogeneously pigmented cells, suggesting heterogeneous capacities for plasticity and clonogenicity amongst HPCs.

**Figure 4.**
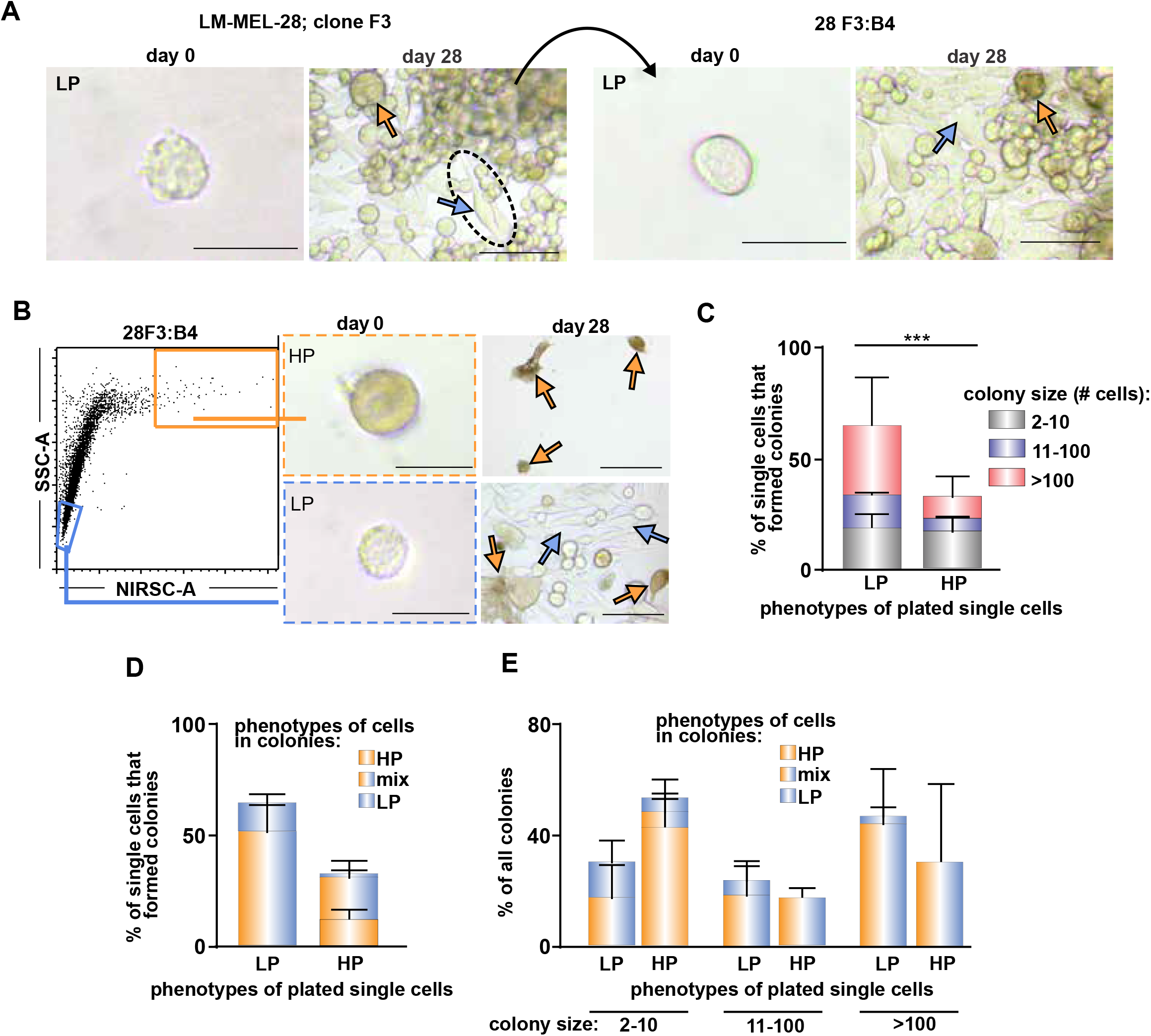
Clonal passageability of LP and HP cells. **A)** Generation of a clonal LM-MEL-28 cell line. A single SSCloNIRSClo LP cell (clone F3) was expanded to generate a colony containing both LP (blue arrows) and HP (orange arrows) cells. A single SSCloNIRSClo LP cell (clone B4) was subsequently expanded from the F3 line to generate a heterogeneously pigmented clonal line (28F3:B4). **B)** Single HP or LP 28F3:B4 cells were sorted based on SSC/NIRSC, assessed for pigment by light microscopy, then cultured for 28 days. Bars: 25μm (single cells) and 100 μm (cultures). **C)** Quantitation of colony (>2 cells) formation and the number of cells per colony. Bars: averages ± SD from 2 independent experiments. n ≥ 32 single cells/phenotype/experiment. ***p<0.001, t-test. **D)** Quantitation of colony phenotypes grown from single LP or HP 28F3:B4 cells. Bars: averages ± SD from 2 independent experiments. n ≥ 32 single cells/phenotype/experiment. **E)** Quantitation of colony phenotypes stratified by size. Bars: averages ± SD from 2 independent experiments. n ≥ 10 (HP) and n ≥ 27 (LP) colonies/experiment.

### Cell surface marker expression in LPCs and HPCs

Surface protein markers have been identified of melanoma cell subpopulations with increased tumorigenic potential, including CD271 (23, 24) and ABCB5 (46, 47), although these observations were not reproduced in highly permissive assays (12, 25, 37). We thus tested the expression of 11 cell markers previously associated with increased melanoma tumorigenicity (CD271, JARID1B, ABCB5, ALDH1A1, c-Kit, CD49f, LICAM, CD51, SOX10, A2B5 and GD2) in 28F3:B4 LPCs and HPCs. By flow cytometry, we were unable to detect notable expression of c-Kit, CD49f, ALDH1A1, L1CAM or CD51 in either LPCs or HPCs (**Figure S5A-E**). Minor populations of cells positive for CD271, JARID1B, ABCB5 (cell surface and intracellular) and SOX10 were detected at similar proportions in both LPCs and HPCs (**Figure 5A-E**). A2B5 and GD2 expression were higher in LPCs than HPCs (61% vs 33% (**Figure 5F**) and 81% vs 58% (**Figure 5G**), respectively). Thus, we found no evidence for enrichment of previously identified cell surface markers in tumorigenic LPCs.

**Figure 5.**
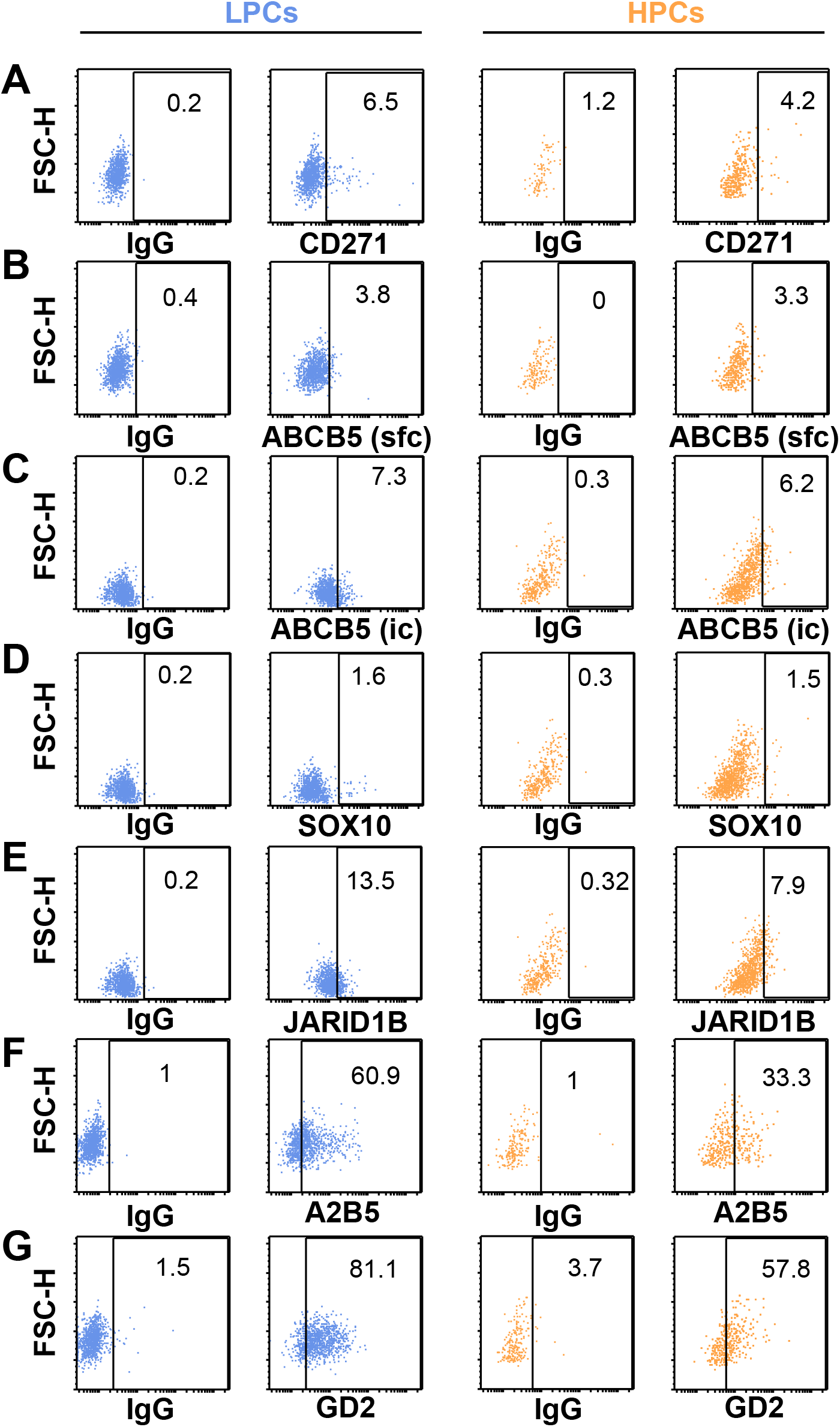
Expression of cell markers in LP and HP melanoma cells. Flow cytometric analysis of LP (blue) or HP (orange) 28F3:B4 cells following staining with IgG controls, or antibodies directed against **A)** CD271, **B)** ABCB5 (surface), **C)** ABCB5 (intracellular), **D)** SOX10, **E)** JARID1B, **F)** A2B5, and **G)** GD2. Background fluorescence was determined separately for LP and HP cells using IgG controls. % of cells indicated in each positive gate.

### Molecular differences between LPCs and HPCs

To identify molecular mechanisms linked to more and less clonogenic/tumorigenic melanoma cell states associated with pigment production, we performed RNAseq on LPCs and HPCs isolated by flow cytometry from B16-F10 and 28:F3:B4 cells. For each cell line, over-representation analysis (39, 40) was performed, and significant enrichment of Gene Ontology (GO) (48) were assigned, based on genes that showed increased expression in each cell phenotype compared to the other. Although melanocytic differentiation genes were strongly upregulated in B16-F10 cells (but surprisingly not in 28:F3:B4 cells; data not shown), upregulated genes in HPCs from both cell lines were overrepresented in the ‘p53’ Hallmark Gene Set and associated with GO terms linked to regulation of cell cycling and cell growth (**Figure 6A**). Upregulated genes in LPCs from both lines were overrepresented in the Hallmark Gene Sets ‘MYC’ and ‘MTORC1’ and associated with ribosome biogenesis and non-coding RNA processing by GO assignment (**Figure 6B**).

**Figure 6.**
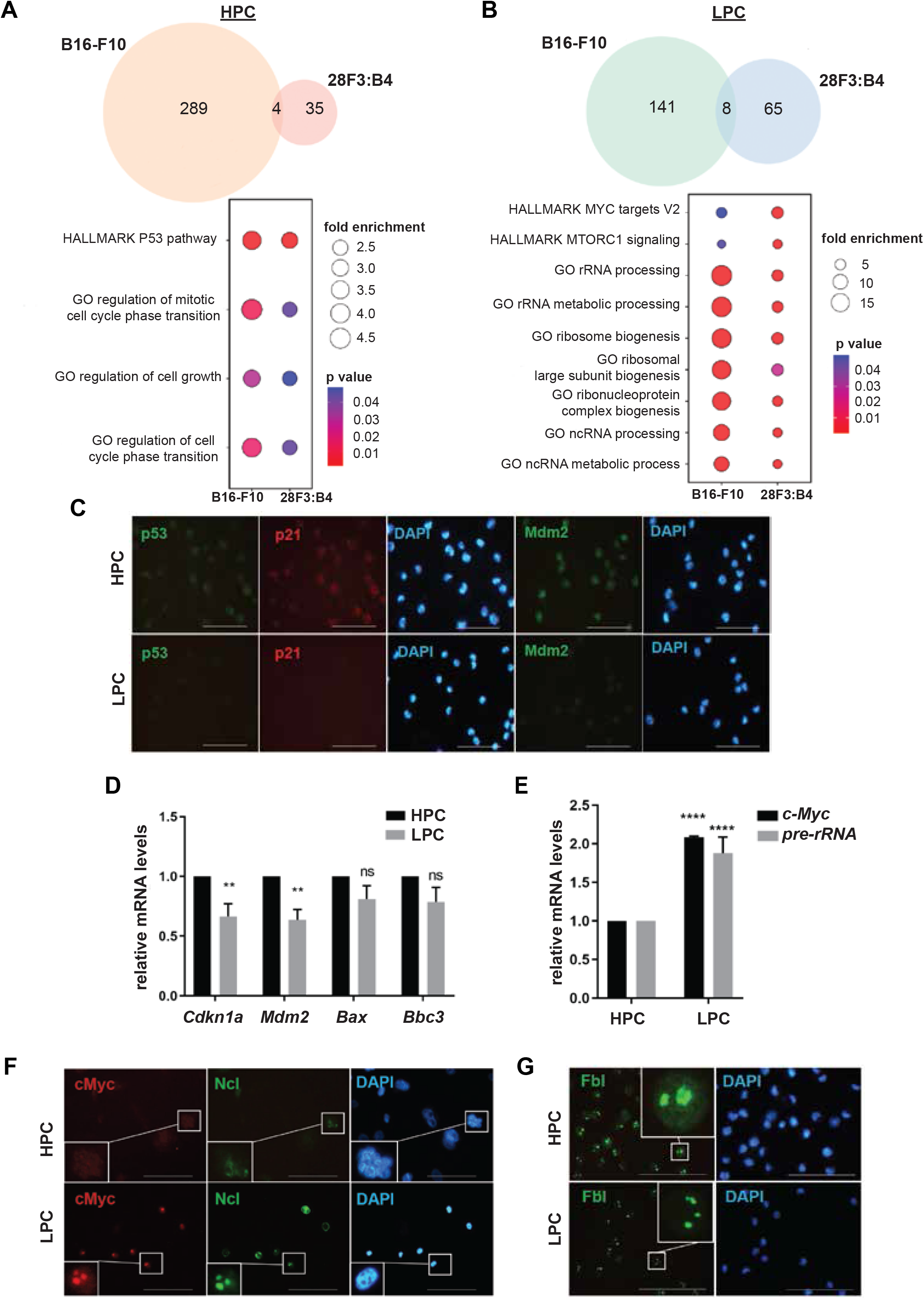
Mechanisms of regulation of LP and HP cells revealed by transcriptional profiling. Gene ontology - biological process (GO-BP) and HALLMARK pathways overrepresented by significantly up-regulated genes in **A)** HP or **B)** LP cells isolated from B16-F10 and 28F3:B4. Venn diagrams (top) enumerate significantly overrepresented pathways, with listed pathways (bottom) common between B16-F10 and 28F3:B4. **C)** HPCs and LPCs from B16-F10 cells were immunostained with antibodies against p53, p21, and Mdm2. DAPI: nuclei. Bars: 100 μm. **D)** p53 target gene (*Cdkn1a, Mdm2, Bax* and *Bbc3*) expression and **E)** *c-Myc* mRNA and *45S* pre-rRNA levels in HPCs and LPCs from B16-F10 cells were measured by qRT-PCR. **p<0.01, ***p<0.001, ****p<0.0001, t-tests. **F)** and **G)** HPCs and LPCs from B16-F10 cells were immunostained with antibodies to c-Myc, Ncl, and Fbl. DAPI: nuclei. Bars: 100 μm. Insets: magnified images of selected cells.

Evidence of p53 activation was also evident at the protein level in purified B16-F10 HPCs, which showed increased expression of p53 itself, as well as p53 targets p21 and Mdm2 (**Figure 6C**), *Cdkn1a, Bax* and *Bbc3* (**Figure 6D**). Similar activation was observed in 28:F3:B4 HPCs (**Figures S6A-B**). Upregulation of c-MYC in LPCs was confirmed by qRT-PCR (**Figure 6E and S6C**), and increased nucleolar localization of nucleolin and fibrillin, as well as increased pre-RNA, were consistent with upregulated ribosome biogenesis in LPCs (**Figures 6E-G and S6C-E**).

### Inhibition of TOP2B promotes non-clonogenic HPC phenotypes

As ribosome biogenesis has a critical role in cancer progression (49, 50), we hypothesized that suppression of this process using CX5461, an inhibitor of topoisomerase 2 beta (TOP2B) in clinical development (51–55), would inhibit LPCs. Treatment with CX5461 of sorted B16:F10 and 28F3:B4 LPCs induced cellular pigmentation (**Figures 7A and S7A**) and higher proportions of SSC^hi^NIRSC^hi^ HPCs by flow cytometry (**Figures 7B and S7B**), relative to vehicle-treated control LPCs. Although CX5461 did not induce substantial apoptosis (**Figures 7C and S7C**), treated LPCs were enlarged and displayed increased X-galactosidase staining compared to controls, consistent with induction of cell senescence (**Figures 7A and S7A**). This senescent state was irreversible, as LPCs pre-treated with CX5461 displayed markedly reduced clonogenicity when re-cultured and passaged without CX5461 (**Figures 7D, 7E, S7D and S7E**). Notably, CX5461-treated LPCs also displayed molecular features of HPCs in delocalized nucleolin, upregulation of p53, and substantially overlapping transcriptional profiles (**Figures 7F, 7G, S7F, S7G and S7H**). These data indicate that TOP2B inhibition induces pigmented, poorly clonogenic, and irreversibly senescent states in melanoma cells.

**Figure 7.**
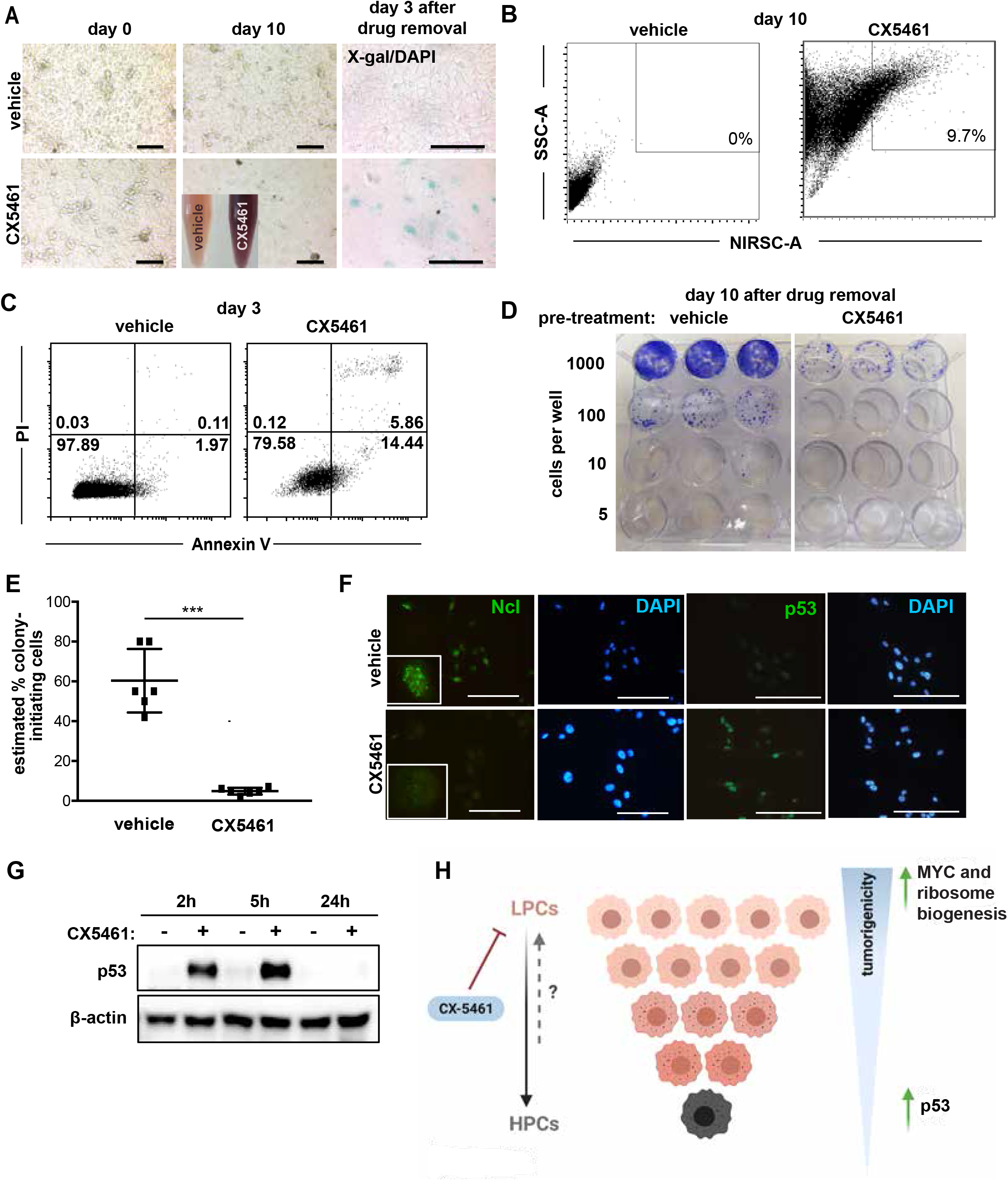
CX5461 treatment promotes melanoma cell pigmentation and loss of tumorigenicity. **A)** Cell pigmentation by light microscopy in purified LP B16-F10 cells exposed to 1μM CX5461 or vehicle control for 3 days. Inset: supernatants from vehicle- (left) and CX5461- (right) treated cultures. X-galactosidase (X-gal) staining after CX5461 or vehicle treatment and then drug withdrawal for 3 days. Bars: 100 μm. **B)** Cell pigmentation was evaluated by flow cytometry of SSC and NIRSC-A 10 days after drug removal. % of SSChiNIRSChi cells are shown. **C)** B16-F10 cells were treated with vehicle or 1 uM CX5461 for 3 days and apoptosis levels evaluated by FACS flow cytometry AnnexinV/ PI labelling. **D)** In vitro clonogenicity of purified LP B16-F10 cells treated with 1μM CX5461 or vehicle for 3 days followed by drug withdrawal for 10 days and then replating in equal numbers for colony assays. **E)** Extreme limiting dilution analysis of the frequency of colony-initiating cells in cultures in each condition in D). ***p<0.001, student t-test. **F)** B16-F10 cells were vehicle or 1 μM CX5461 treated for 24h and immunostained with antibodies Ncl and p53. DAPI: nuclei. Bars: 100 μm. Insets: magnified images of selected cells. **G)** B16-F10 cells were vehicle or 1 μM CX5461 treated for 2h, 5h or 24h and lysates immunoblotted with anti-p53 antibody. β-Actin: loading control. **H)** Proposed model of ‘inverted pyramid’ hierarchical organisation of LPCs and HPCs within pigmented melanomas. Supported by activated MYC and ribosome biogenesis, abundant LPCs proliferate to generate new LPCs that maintain tumorigenic potential and also other progeny that acquire features of melanocytic differentiation. Although many of the latter cells may retain an ability plastically to re-generate LPCs and also intermediately pigmented cells, some progenies are highly pigmented and irreversibly lose tumorigenic potential in association with p53 upregulation.

## DISCUSSION

We report a novel, flow cytometry-based method for prospectively isolating viable cells according to their content of melanin pigment, using NIRSC and SSC as key parameters to distinguish HPC and LPCs. This permitted prospective in vitro and in vivo analyses of the biological consequences of production of melanin pigment in melanoma cells, revealing striking differences in the abilities of cells with low versus high pigment levels to propagate disease and a hierarchy of melanoma cell development within cell lines and tumors (Figure 7H). The reduced clonogenic and tumorigenic potential of HPCs was associated with up-regulation of P53 and perturbation of cell cycle regulators, whereas highly tumorigenic LPCs displayed MYC activation and increased ribosome biogenesis (Figure 7H). Importantly, the latter was exploitable therapeutically, wherein TOP2B targeting promoted senescent, HPC-like phenotypes in LPCs, and irreversible loss of clonogenicity.

Unlike intratumoral cellular hierarchies in classical pyramidal models of cancer progression (56) in which less- or non-tumorigenic cells are more abundant than the highly tumorigenic cells that beget them, we found an inverted structure in melanoma. Consistent with previous observations that remarkably high proportions of melanoma cells harbor intrinsic tumorigenic potential (57), LPCs were vastly more abundant than HPCs in all melanomas we evaluated with heterogeneously pigmented cells. Moreover, LPCs typically gave rise to heterogeneously pigmented colonies in clonal culture, and also frequently generated large LPC-only colonies, indicating their ability to propagate the LPC phenotype, as would be required in an ‘inverted pyramid’ cellular model of the disease (Figure 7H).

In contrast, a substantial proportion of single HPCs generated small HPC-only colonies, and markedly reduced or absent ability was found in HPCs to form tumors and propagate disease, even in highly permissive tumorigenesis assays (57). Indeed, despite exhaustive testing of numerous markers including CD271 (12), a high level of melanin pigment production is the first marker we have identified in melanoma cells that reliably predicts loss of tumorigenic potential.

Plastic relationships between LPCs and HPC were occasionally but notably observed in some assays. The proportion of these might have been underestimated in our study, which only evaluated cells at extremes of the spectrum of intracellular pigmentation as we sought to derive maximally pure populations of LPCs and HPCs for biological and molecular assays. However, substantial populations of cells with medium and low levels of melanin pigment (mixed with phenotypic LPCs) resided between these extremes. Pigmentation differences between these populations were difficult to resolve with NIRSC-based flow cytometry, and thus not extensively evaluated. We therefore did not test prospectively whether pervasive plastic relationships exist between LPCs and melanoma cells with at least low to moderate levels of pigment, as predicted in the ‘phenotype switching’ model of disease propagation (58).

Although our data, therefore, are not in conflict with the ‘phenotype switching’ model, they do nuance this model by demonstrating that some transitions to the HPC state are irreversible and characterized by loss of clonogenic and tumorigenic potential. Such transitions might be infrequent in melanomas, but their existence *per se* raises the question as to why high levels of melanin pigment might be linked to decreased malignant behavior in melanoma cells. On one hand, HPC states might represent endpoints of developmental programs of melanocytic differentiation activated in the progeny of some LPCs that generate fully differentiated, post-mitotic cell states. In this case, upregulation of melanocyte differentiation genes would be expected in HPCs. Although this was observed in one cell line, it was not a ubiquitous feature. Rather, dysregulation of P53 and of cell cycle genes were consistently prominent in HPCs.

Alternatively, melanin production might be induced stochastically or as a result of environmental or cell intrinsic stresses in tumors. Anecdotally, melanin deposition is often more prominent in patient melanomas at margins between viable and non-viable regions within tumors, consistent with the notion that micro-environmental factors can influence melanin production in melanoma cells. As melanin pigment can itself be toxic to cells (59–61), associated with DNA damage (62), a mechanism of spontaneous generation of HPCs is suggested wherein melanin production in melanoma cells is induced cell extrinsically to a point from which a cell intrinsic, positive feedback loop is established of increasing melanin deposition and genotoxic stress that ends in functional senescence.

Our finding of spontaneous development of irreversible loss of clonogenicity/tumorigenicity in HPCs raises the intriguing possibility that HPC phenotypes could be promoted therapeutically to inhibit melanoma progression. Towards this, as our gene expression profiling data indicated activation in LPCs of ribosome biogenesis, we tested pharmacological treatment with CX-5461, an agent in clinical development (55) and with promising activity against a range of cancers (51, 52, 55, 63, 64) that is mediated by inducing DNA damage through TOP2B interactions (65) and inhibiting ribosomal RNA (52). Although CX-5461 induced some cell death, its dominant anti-melanoma effect was induction of senescent HPC phenotypes and transcriptomes. These findings provide support not only for clinical development in melanoma of TOP2B targeting, but also for the notion that melanin deposition in melanoma cells both potentiates and results from genotoxicity.

Our discovery and exploitation of the NIR scattering properties of melanin, and resultant development of a novel flow cytometric approach to analysis of cells in the melanocytic lineage, provides an unprecedented opportunity to functionally characterise these cells based on a classical phenotypic marker of melanocyte differentiation. This will be useful not only in melanoma research, as we demonstrate here by identifying mechanisms through which melanoma cells spontaneously become and may be rendered incapable of disease propagation, but also to advance understanding of normal and pathological melanocyte development.

## ACKNOWLEDGEMENTS

This work was supported by the Melanoma Research Alliance (USA), the National Health and Medical Research Council of Australia, Melanoma Research Victoria, and the Victorian Cancer Biobank. M.S. was supported by Fellowships from Pfizer Australia, veski, and the Victorian Cancer Agency. We thank staff of the flow cytometry and experimental animal facilities of the Peter MacCallum Cancer Centre and Monash University’s Central Clinical School.

## SUPPLEMENTARY FIGURE LEGENDS

**Figure S1. LPC in tumors are melanoma cells (relating to Figure 1) A)** Representative histological images of pigmented patient melanomas stained with the melanin stain, Schmorl’s **B)** Adjacent sections of human melanomas stained for the melanin stain, Schmorl’s, and S100B as a marker of melanoma

**Figure S2. Detection of endogenous pigment by flow cytometry (relating to Figure 2)** Flow cytometric detection of signal following excitation of amelanotic A375 or heterogeneously melanotic B16-F10 cells with near-infrared (792nm) wavelengths and signal detection with filters that either capture (710, 745, 750 LP or 817/25 BP) or eliminate (800, 900 LP) the excitation wavelength

**Figure S3. Flow cytometry gating strategies (relating to Figure 2) A)** Flow cytometric analysis of SSC and NIRSC in amelanotic LOX-IMV-1 cells or B16-F10 cells treated for 72 hours with 20mM Forskolin (Forsk) + 100mM IBMX, or control vehicle. % of SSC^hi^NIRSC^hi^ cells is indicated. Inset images show FM staining for melanin and % FM+ve cells. **B)** Fluorescence-activated cell sorting (FACS) of heterogeneously pigmented cells from a pigmented PDX melanoma from patient M176 based on SSC and NIRSC. The proportions of HPC cells, as determined by FM staining, is indicated for each gate. Scalebars = 50um.

**Figure S4. LP cells have increased clonogenic and tumorigenic potentials (relating to Figure 3) A)** LP and HP melanoma cells were sorted based on SSC/NIRSC from a pigmented PDXs from patient M840 and M172 (photos inset) and injected subcutaneously into NOD-scid IL2Rγ^null^ (NSG) mice at limiting numbers (10-1000 cells). Images show FM staining in sorted HP and LP populations. Scalebars = 50μm **B)** Tumor latencies and **C)** growth rates of tumors grown from either purified LP or HP cells from 2 patient PDXs *p< 0.05 **p< 0.005. **D)** Bioluminescent detection of metastases in organs of mice bearing tumors grown from either LP or HP cells from patient M840, and quantitation of mice with macroscopically detectable metastases bearing tumors grown from either LP or HP cells from 2 PDXs. n = 5-6 mice/phenotype. Lu: Lung; He: Heart; Sp: Spleen; Li: Liver; Rep: Reproductive organs; Ki: Kidneys; Br: Brain; GI: Gastrointestinal tract; St: Stomach. **E)** Single LP or HP B16-F10 cells were sorted by flow cytometry (1 cell/well). At day 0 each cell was assessed for the presence or absence of pigment by light microscopy (upper panel), then allowed to proliferate for 21 days in culture. At experimental endpoint the pigment phenotypes of resultant colonies were assessed by light microscopy for the presence of LP (blue arrows) and/or HP (orange arrows) cells (lower panel). Scalebars = 25μm (upper) and 100μm (lower). **F)** Quantitation of colony phenotypes grown from single LP or HP cells. Bars represent the averages ± SD from 2 independent experiments. n ≥ 17 single cells/phenotype/experiment. **G)** Quantitation of colony phenotypes stratified by size. Bars show averages ± SD from 2 independent experiments. n ≥ 7 (HP) and n = 32 (LP) colonies/experiment. ND=colony phenotype not determined. **H)** Single LP 28F3:B4 cells were sorted by flow cytometry (1 cell/well). At day 0 each cell was assessed for the presence or absence of pigment by light microscopy, then allowed to proliferate for 32 and 63 days in culture. At experimental endpoint the pigment phenotypes of resultant colonies were assessed by light microscopy.

**Figure S5. Expression of putative CSC markers in LP and HP melanoma cells (relating to Figure 5)** Flow cytometric analysis of LP (blue) or HP (orange) 28F3:B4 cells following staining with IgG controls, or antibodies directed against **A)** c-Kit, **B)** CD49f, **C)** ALDH1A1, **D)** CD51, **E)** L1CAM. Background fluorescence was determined separately for LP and HP cells using IgG controls. % of cells in each positive gate is indicated.

**Figure S6. Characterisation of HP and LP 28F3:B4 cells. A)** Immunostaining of HP and LP 28F3:B4 cells with antibodies to p53, p21, and MDM2. DAPI: nuclei. Bars: 100 μm). **B)** p53 target gene (*CDKN1A, MDM2, BAX* and *BBC3*) expression in HP and LP 28F3:B4 cells was measured by qRT-PCR. *p<0.05, **p<0.01 and ****p<0.0001, student t-test. **C)** *C-MYC* mRNA and *45S* pre-rRNA levels were assessed by qRT-PCR in HP and LP 28F3:B4 cells. ***p< 0.001, student t-test. **D)** HP and LP 28F3:B4 cells were immunostained with antibodies to c-MYC, NCL, and **E)** FBL. DAPI: nuclei. Bars: 100 μm. Insets: magnified images of selected cells.

**Figure S7. CX5461 treatment promotes melanoma cell differentiation (relating to Figure 7).** Purified LP 28F3:B4 cells exposed to 1 μM CX5461 or vehicle control for 10 days. **A)** Assessment of cell pigmentation by light microscopy was performed in addition to X-galactosidase staining of purified LP cells exposed to 1μM CX5461 or vehicle control for 10 days, followed by drug withdrawal for 5 days. Scale bars=100μm. **B)** Pigmentation levels were evaluated by FACS analysis of SSC and NIRSC-A profiles. % SSChiNIRSChi is indicated. **C)** 28F3:B4 cells were either vehicle or 1 μM CX5461 treated for 3 days, and apoptosis levels were evaluated by FACS after AnnexinV/ PI staining. **D)** In vitro clonogenicity of purified LP 28F3:B4 cells exposed to 1μM CX5461 or vehicle control for 10 days followed by drug withdrawal for 28 days. **E)** Extreme limiting dilution analysis to estimate % colony-initiating cells in 28F3:B4 LP cells exposed to either CX5461 or vehicle control. ***p< 0.001, student t-test. **F)** 28F3:B4 cells were either vehicle or 1 μM CX5461 treated for 24h and immunostained with NCL (green), and p53 (green). DAPI was used as the nuclear marker. (Scale bar, 100 μm). Insets show magnified images of the selected cell. **G)** 28F3:B4 cells were either vehicle or 1 μM CX5461 treated for 2h, 5h and 24h and lysates were immunoblotted with anti-p53 antibody. β-Actin antibody was used as the loading control. **H)** Heat map of differentially expressed genes up and downregulated in 1 μM CX5461 treated B16-F10 cells and HP B16-F10 cells. Colour scale represents the log fold change compared to parental controls.

## Notes

### Competing Interest Statement

The authors have declared no competing interest.

