## Supplementary figures and images for "Prospective isolation according to melanin pigment content of melanoma cells with heterogeneous potentials for disease propagation"

### Supplemental Figures

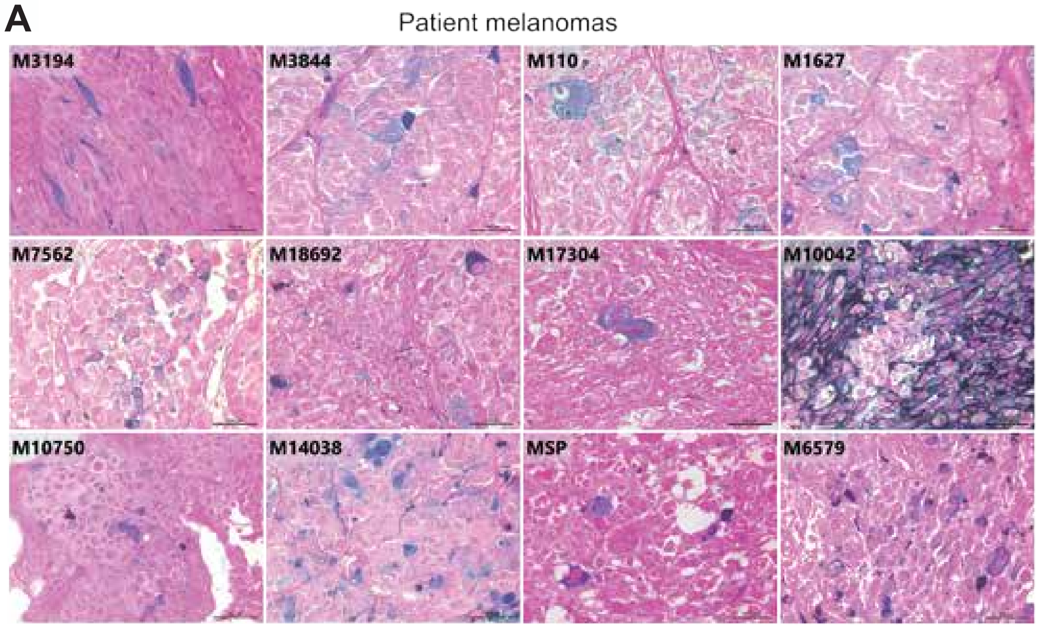

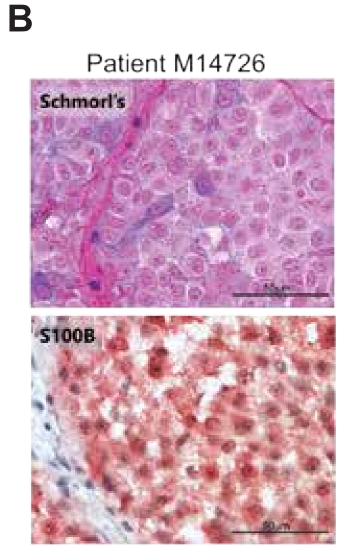

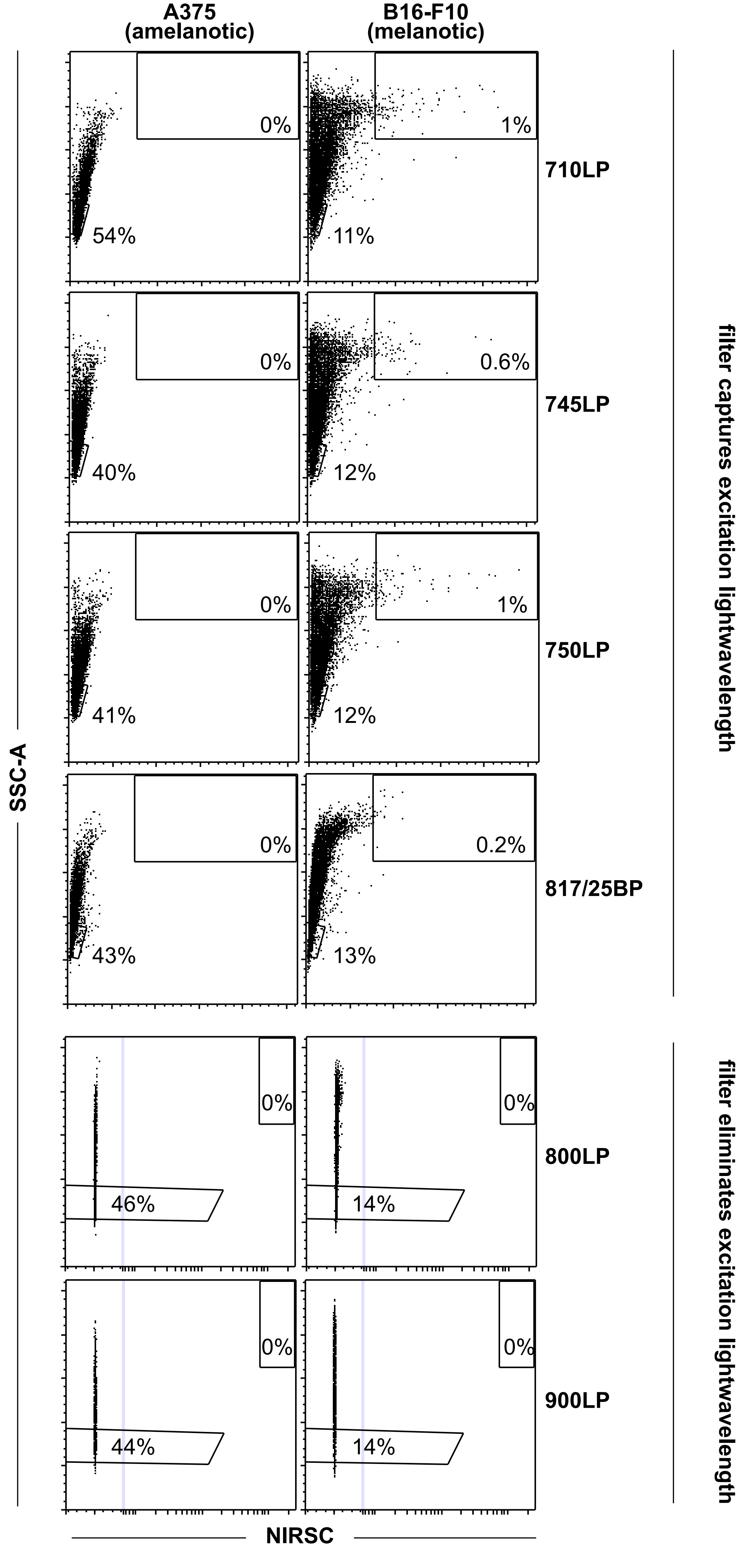

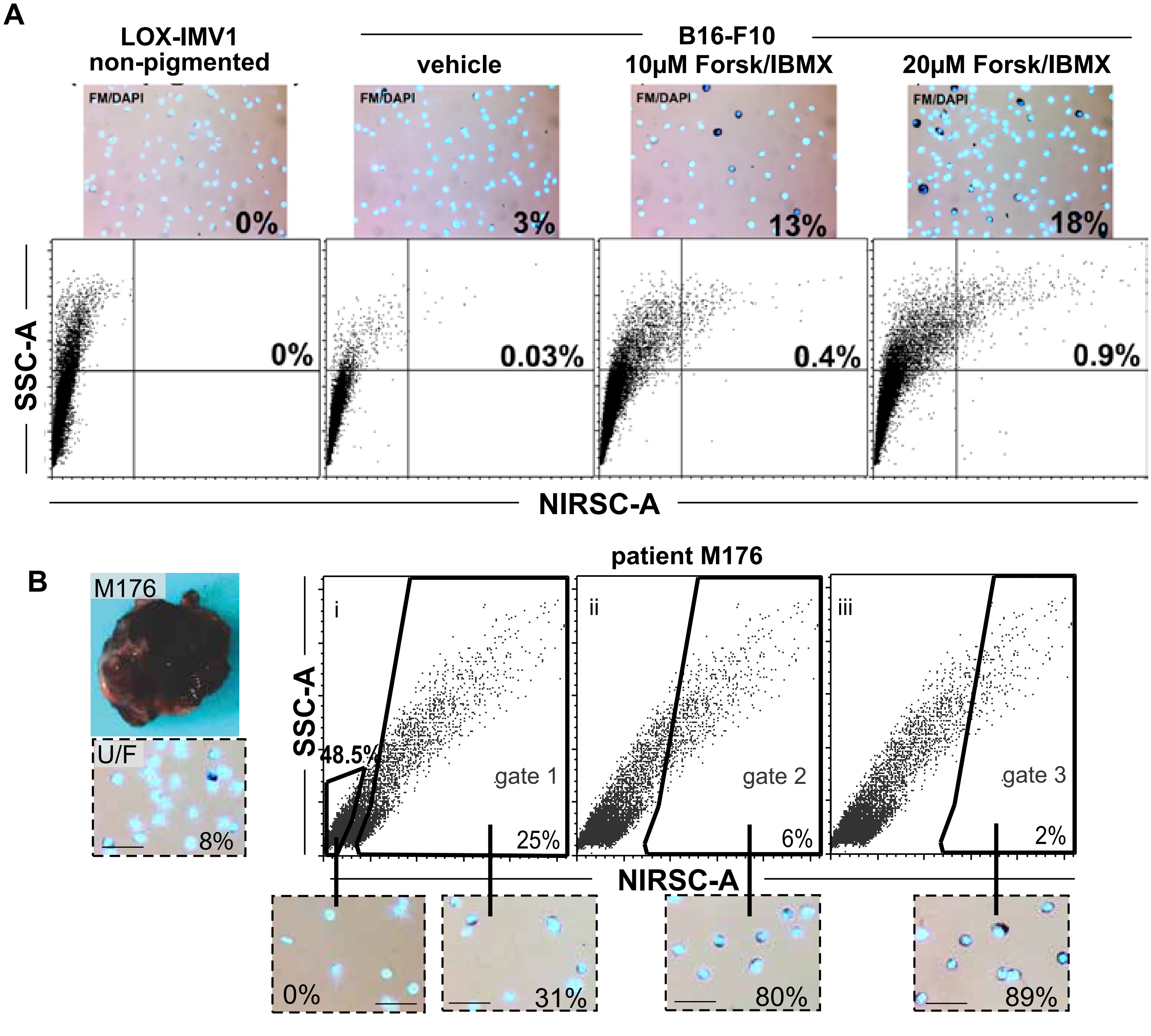

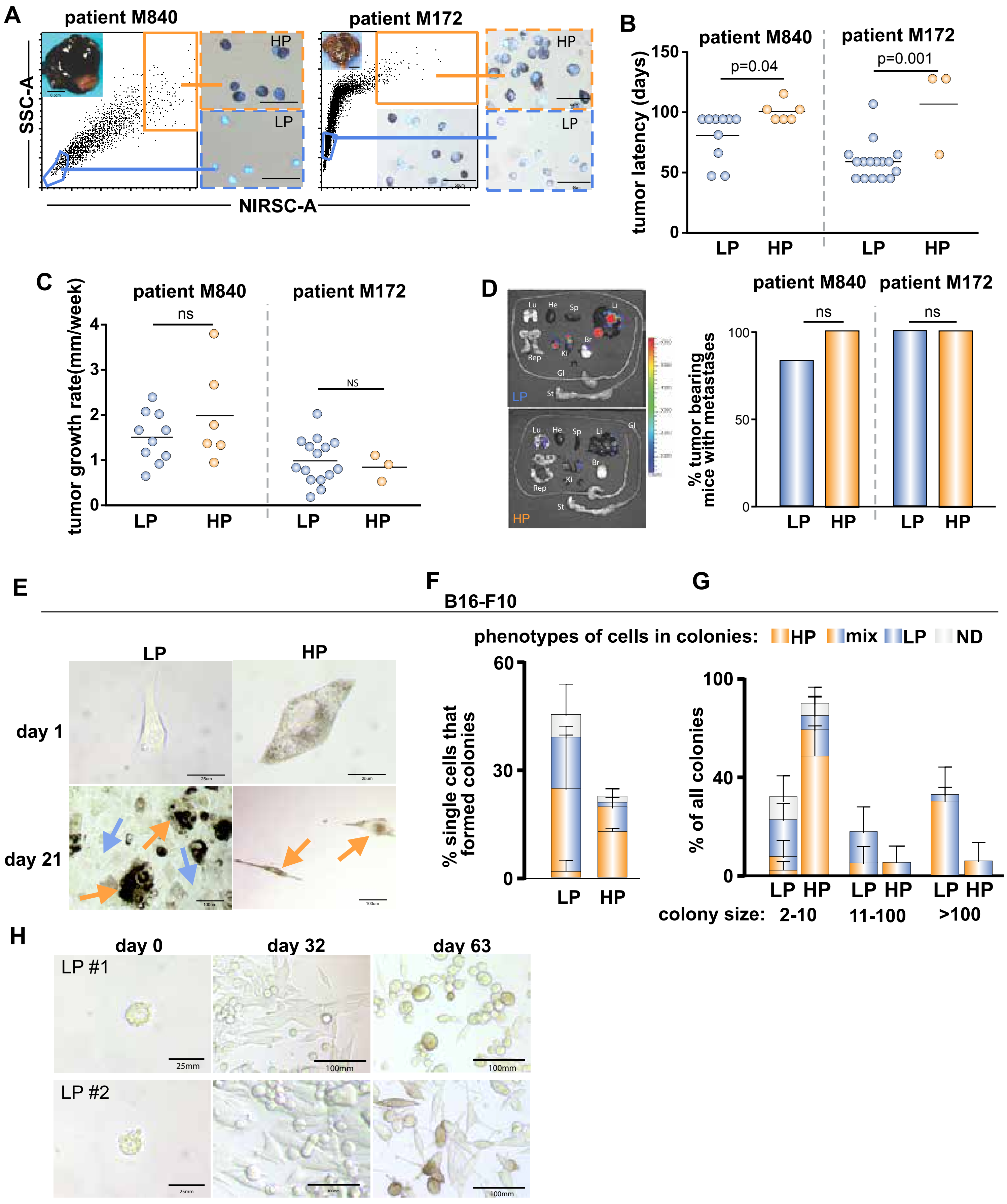

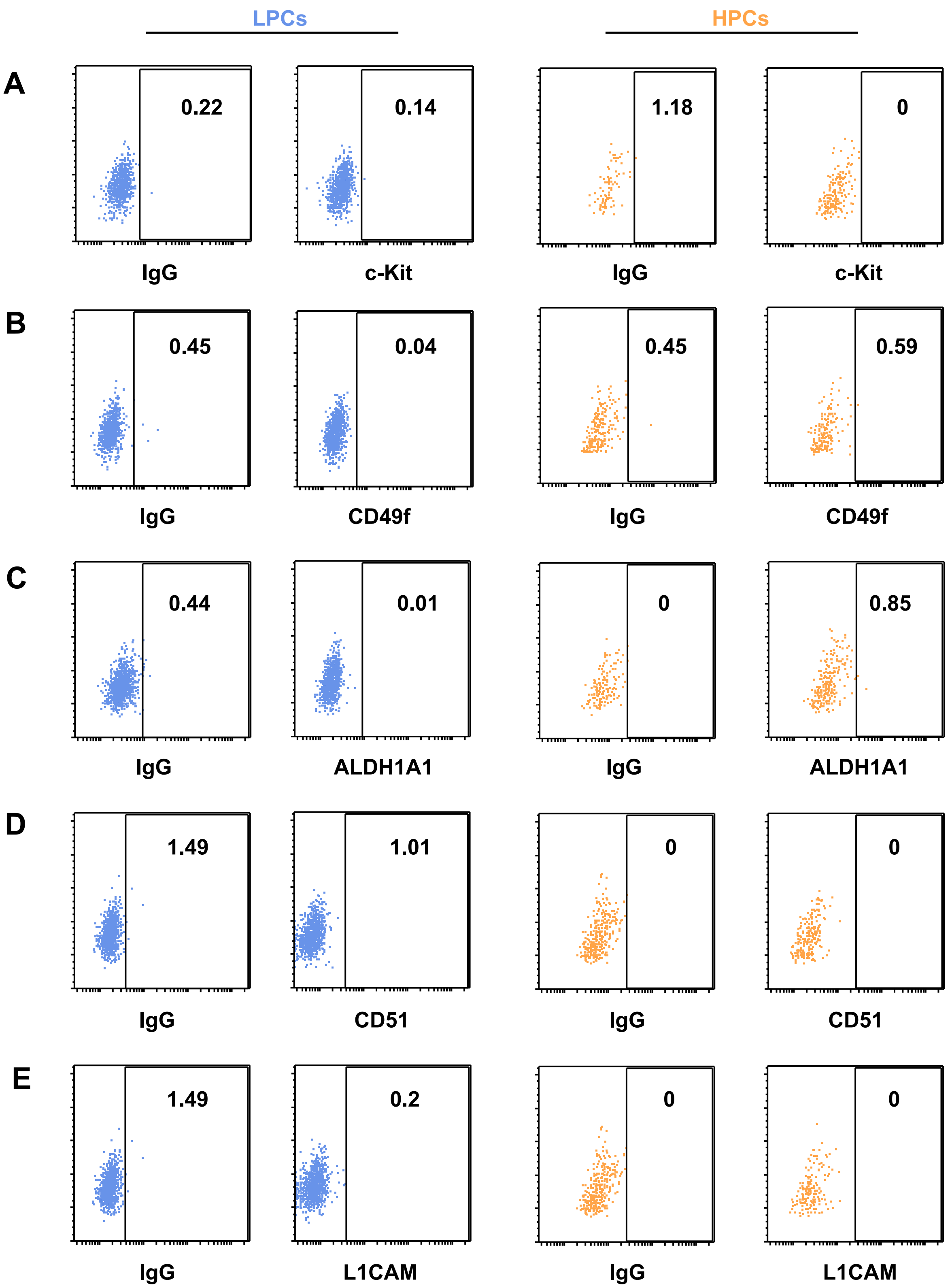

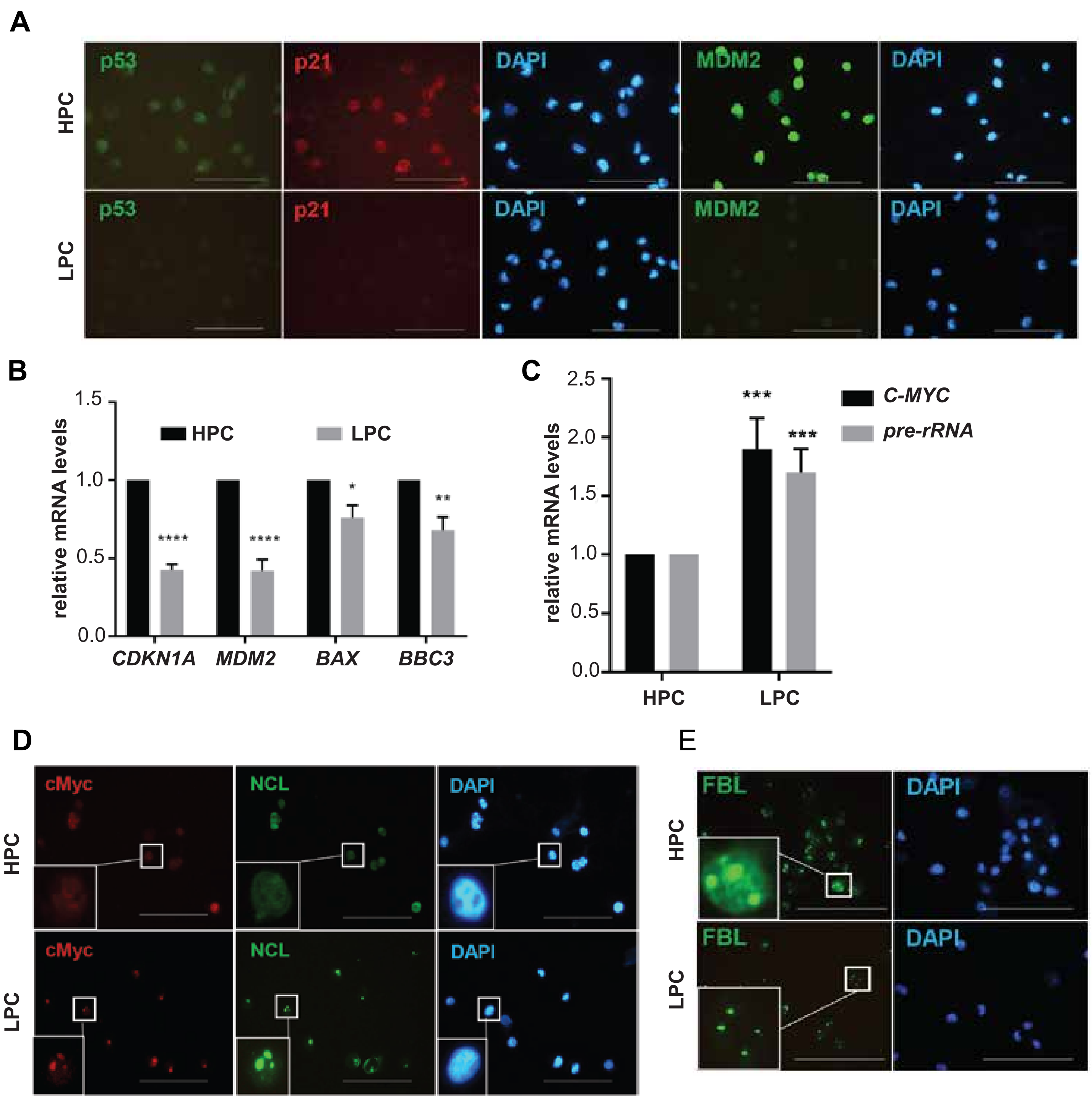

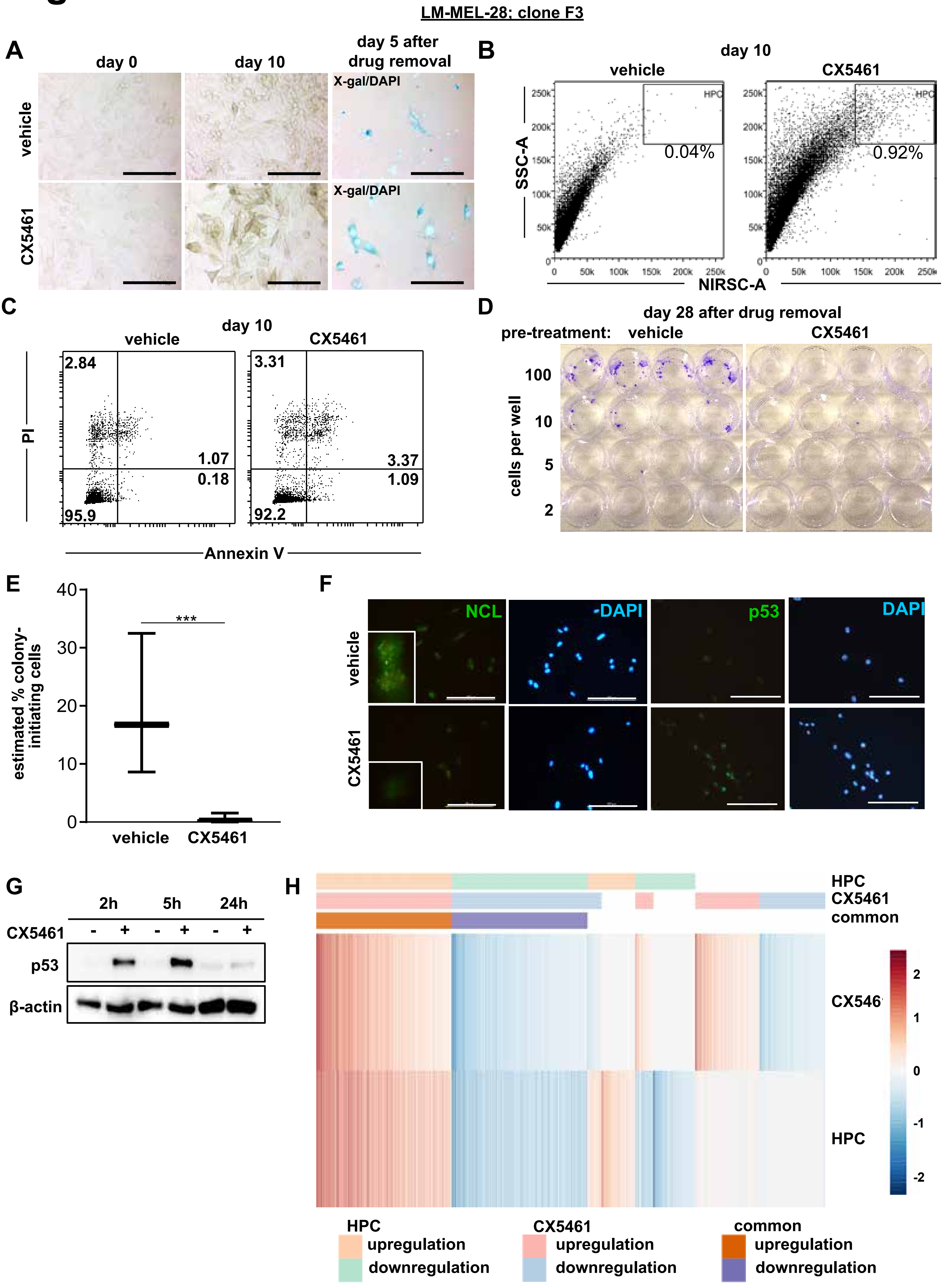
